# 3D Printed Skull Cap and Benchtop Fabricated Microwire-based Microelectrode Array for Custom Rat Brain Recordings

**DOI:** 10.1101/2022.09.26.509528

**Authors:** Dongyang Yi, Jeremiah P. Hartner, Brian S. Ung, Harrison L. Zhu, Brendon O. Watson, Lei Chen

## Abstract

Microwire microelectrode arrays (MEAs) have been a popular low-cost tool for chronic electrophysiological recordings. Multi-MEA implantations can reveal electrical dynamics crucial to brain function. However, both the fabrication and implantation procedures for multi-MEAs on a single rodent are time-consuming and highly manual skill-dependent for quality. To enable in-house design, fabrication, and implantation of custom microwire MEAs, we developed (1) a computer-aided designed and 3D printed skull cap for pre-determined implantation locations of each MEA and (2) a benchtop fabrication approach for low-cost custom microwire MEAs. A proof-of-concept design of 32-channel 4-MEA (8-wire each) recording system was prototyped and tested through Sprague Dawley rat recordings. The skull cap design based on CT-scan of single rat conforms well with multiple Sprague Dawley rats of various size, age, and weight with minimal bregma alignment error. The prototyped 32-channel system were able to record spiking activities over 5 months. In comparison with conventional stereotactic surgeries, the skull cap system simplifies the implantation location alignment for each MEA by embedding them into the pre-printed designs, thus dramatically reducing the surgical time and effort and increasing the accuracy and repeatability. Compared to commercially available custom microwire MEAs, this in-house fabrication method enables neuroscience labs to create a custom recording apparatus with lower cost and shorter lead time for design modifications. A new methodology for neuroscience labs to fabricate and insert custom microwire MEAs has been developed and it could be easily generalized to enable low-cost highly-custom multi-region recording/stimulation studies.

**Highlights:**

- 3D printed skull caps as implantation platform for multi-region rat brain recording
- Computer-aided design for custom cap geometry and predetermined implantation location
- Benchtop microwire fabrication approach for in-house custom microelectrode arrays
- Duplicable and generalizable design, fabrication, and implantation methodologies

## 1. Introduction

Electrophysiological recording of single neuron activity is critical for understanding the brain functioning mechanisms [1, 2]. To gain further insight into the dynamic neural mechanisms and interactions across different brain regions, simultaneous chronic recording from multiple locations at high temporal resolution in freely behaving animals is needed [3-5]. The implantation of microelectrode arrays (MEAs) has evolved for decades and has been proven to be a powerful tool to provide high temporal resolution recordings of action potentials [6-9]. Despite the emerging development of silicon based MEAs in recent decades, microwire MEAs, first developed in the 1950s [10], are still a popular choice for neuroscientists due to their low cost [11], high temporal resolution, and great chronic recording performance thanks to their high flexibility and small size [12, 13].

Since the brain is composed of multiple interacting regions, multiple MEAs implanted across regions can reveal fast timescale electrical dynamics crucial to brain function. Current approaches for multiple microwire MEA recordings in rodents rely on repeated single implant craniotomes and other tedious stereotactic-based surgical procedures that are not automated, nor do they take advantage of modern mechanical tools. Specifically, in typical surgeries, implant locations are individually determined within surgery and a separate craniotomy and durotomy are performed at each location [14, 15]. During each implantation, the insertion depth (Dorsal/Ventral) is also controlled by manual operation and measurement through the stereotactic device or linear stage [16, 17]. Temporary apparatus like thin metal bars and 3D printed custom fixtures are used to hold the MEA in place until cured dental cement fixed the electrode at its final position [18, 19]. As a result, three-dimensional positioning accuracy and repeatability of each implantation position are highly dependent on surgical skill and practice. Manual manipulation and alignment of multiple MEAs with a small area of the animal brain surface is labor-intensive and easily leads to complex surgeries lasting many hours. New approaches are needed to simplify the surgery procedure and increase the accuracy and repeatability of multi-MEA insertion locations. Moving towards to use of computer-based designs can open the door to later-future automation and higher throughput. But an initial movement away from all-manual approaches is needed first.

Fabricating MEAs themselves is also a highly manual skill-dependent process. The general fabrication steps involve electrode alignment which counts on manual work with grids and hands-on soldering small wire soldering [20–22], and both procedures are time-consuming. Such a labor-intensive fabrication process has driven many neuroscientists toward commercially available microwire MEAs [23, 24]. These arrays, while ready off-the-shelf, require recording experiments to be designed around the available configurations [16, 25] and would result in higher costs and a longer lead time for customized MEA geometry needs.

In this study, to enable rodent neuroscience groups to design, fabricate, and implant custom microwire MEAs in house, we developed (1) a digitally designed and 3D printed custom skull cap with predetermined insertion locations and depths for accurate but simplified multi-region recordings and (2) a benchtop fabrication approach for low-cost, highly customizable, and in-house made microwire-based MEAs using easily accessible materials and tools with minimal training needed. To demonstrate the developed methodology, a proof-of-concept surgery was planned and conducted to implant four 8-channel planar microelectrode arrays of 50 μm diameter tungsten microwires and 1 mm pitch to broadly record from cortical areas of a Sprague Dawley rat brain.

## 2. Materials and Methods

In this section, the design and fabrication approach for our 32-channel 4-MEA recording system is elaborated as a demonstration of our 3D printed skull cap and benchtop microwire MEA fabrication method. Generalization strategy to move beyond this application-specific design is described at the end of each subsection.

### 2.1. Recording apparatus overview and printed circuit boards design

Figure 1 shows the overview of the recording apparatus proposed for the multiple MEA recording paradigm. Three major components were needed to complete the apparatus: (1) a digitally designed and 3D printed skull cap with structures to guide and position each MEA implantation (elaborated in Section 2.2), (2) assembled microwire-based MEAs with custom configurations (to be described in Section 2.3), and (3) corresponding printed circuit boards (PCBs) for both electrical connection of microwires in each array and connection of multiple arrays to the proposed headstage.

**Fig. 1.**
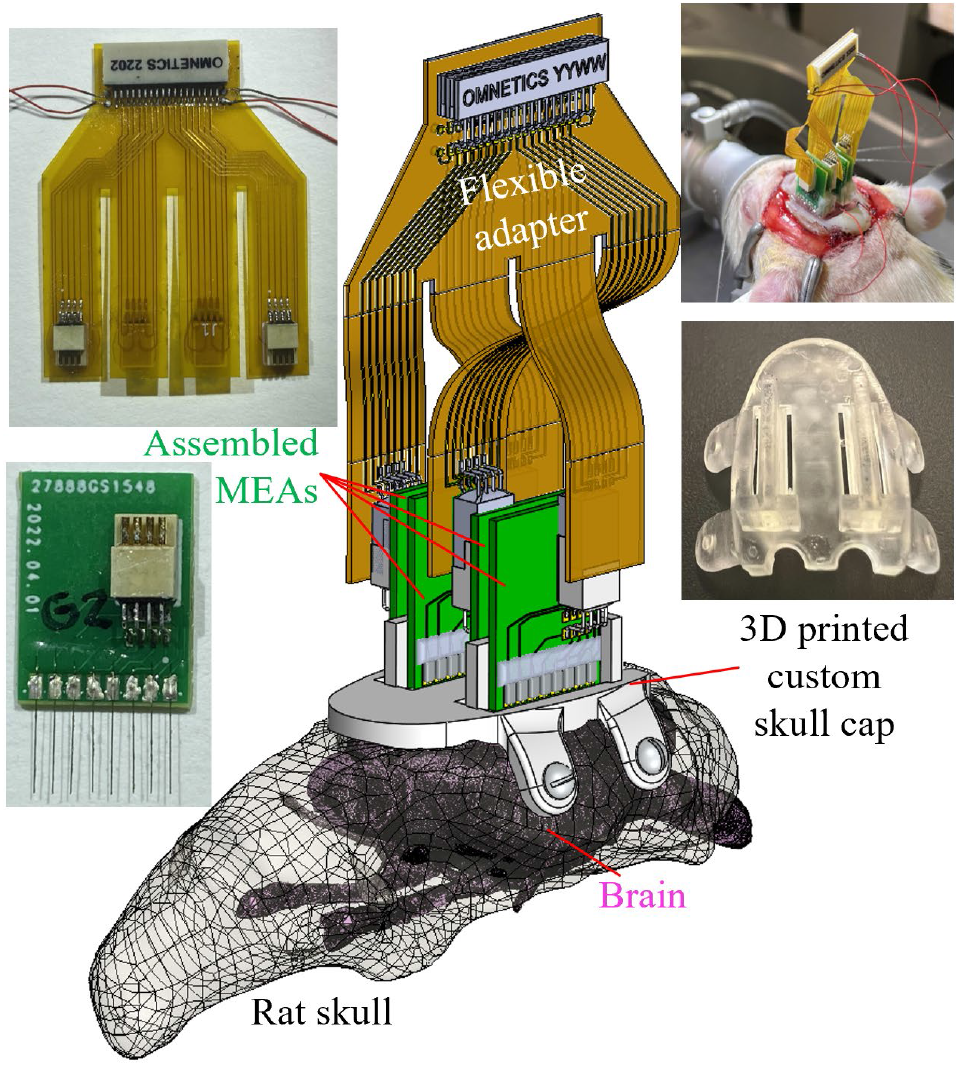
Major components overview of the recording apparatus.

In this demonstration study, the 32-channel recording headstage (RHD 32 by Intan Technologies, Los Angeles, CA) determined the end female connector (A79025 by Omnetics Connector Corporation, Minneapolis, MN) that the developed electrical connection system would be connected to. To provide connections between each of the 32 microwires to one of the pins of the headstage connector, a two-stage PCB system was proposed and developed, as shown in Fig. 2. The first stage (Fig. 2(a)) was a rigid PCB connecting eight (8) microwires in an array to an 8-pin connector (A79613-001 by Omnetics Connector Corporation, Minneapolis, MN). The eight (8) gold pads towards the bottom edge matched the microwire array pitch (1 mm) and would connect to individual microwires. The other eight connection pads were patterned based on the 8-pin Omnetics connector. Note that the connector was placed off-centered on the PCB, which was meant to maximize the rat skull space usage by staggering PCBs together (to be described in Section 2.3 and Fig. 5I). The second stage of the PCB system (Fig. 2(b)) was a flexible PCB adapter connecting four 8-pin connectors from MEA arrays to a male 36 position connector (A79024-001 by Omnetics Connector Corporation, Minneapolis, MN) paired with the 32-channel headstage. Besides 32 microwire connections, the remaining four pins were used for ground and reference wires, as shown in the assembled electronics in Fig. 2(c). To minimize the flexible adapter PCB size, a dual-side circuit board was designed and fabricated with connections to two MEAs on each side. Four long flexible “legs” of the PCB enabled flexible connection after MEA implantation, to be shown in Section 2.4 and Fig. 6(f). Both PCBs were designed digitally (EAGLE by Autodesk Inc., San Rafael, CA) and outsourced for fabrication(PCBminions Inc., Princeton, NJ). The rigid PCB was fabricated with FR-4 glass-reinforced epoxy laminate material, 0.6 mm board thickness, hot air solder leveling surface finish, and 1 oz finished copper. The 2-layer adapter PCB in this study was made of 0.5 mil thick polyimide for optimized flexibility.

**Fig. 2.**
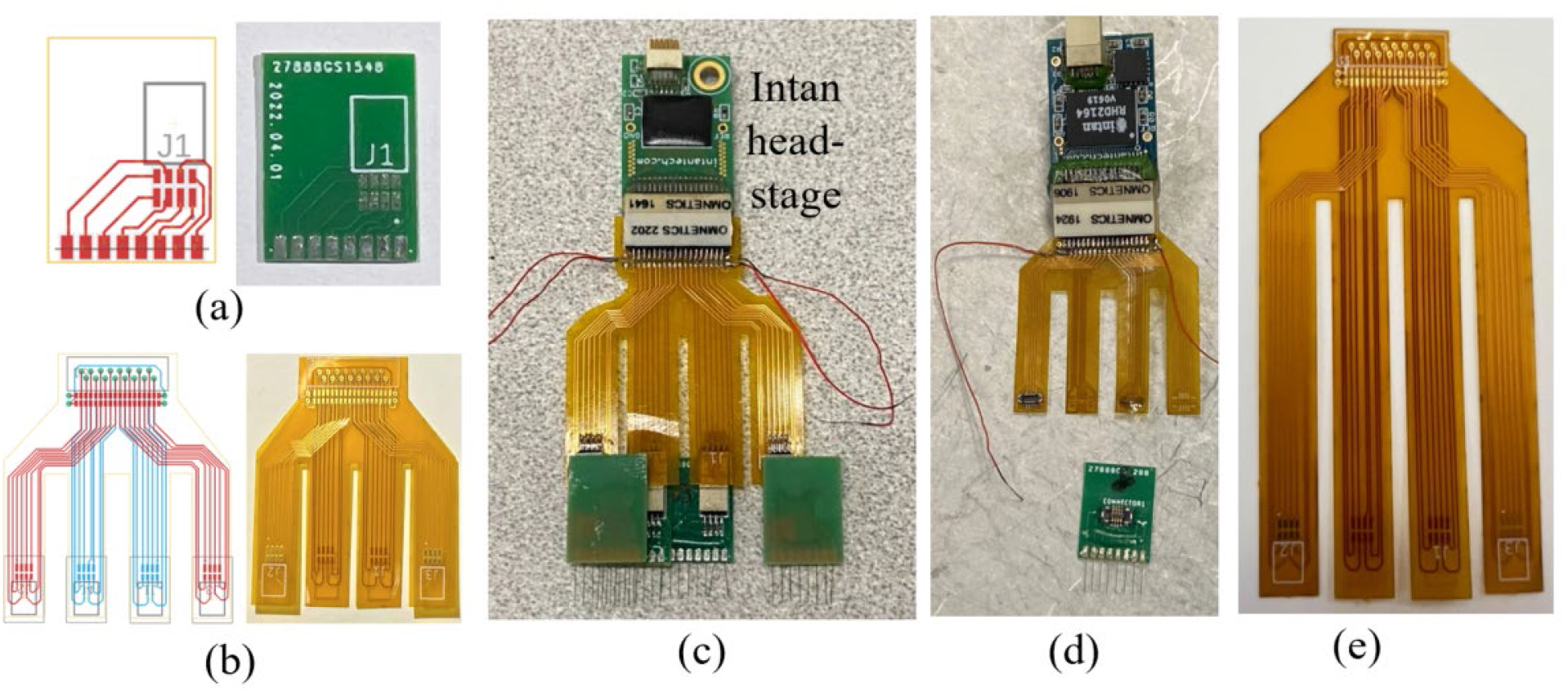
Digital design and fabricated pieces of (a) the microwire MEA PCB and (b) the flexible adapter PCB and (c) the final assembled electronics with headstage. The same design concept could also be implemented with other preferred (d) connectors and/I(e) configurations.

The PCB designs and parameters shown in this proof-of-concept trial were tailored based on the specific recording species and task in this study. The same concept of a two-stage connection system composed of planar rigid and flexible PCBs could be implemented with different MEA pitch sizes, various total channel count and distribution (determining wire number in each MEA), preferred type of connectors (prototype based on Hirose^TM^ connectors shown in Fig. 2(d)), and custom flexible adapter configurations (adapter PCB with longer “feet” shown in I. 2(e)) for specific species and desired recording locations.

### 2.2. Design and fabrication of the 3D printed skull cap

To eliminate the labor-intensive manual alignment and temporary fixation of each MEA during multi-region surgery, this study used a digitally designed and 3D printed skull cap. A computed tomography (CT) scan of the rat skull (Fig. 3(a)) was first conducted using a microCT system (Skyscan 1176 by Bruker Corporation, Billerica, MA) with 35 μm resolution. ImageJ was used to process the microCT images and reconstruct the three-dimensional skull geometry (Fig. 3(b)). The skull geometry was then imported into a computer-aided design (CAD) software (SolidWorks by Dassault Systems Inc., Waltham, MA) for further designs. For a basic solid cap design (Fig. 3(c)), the bottom of the cap matched the top surface of the skull and the top of the cap was designed to be a flat surface as the implantation apparatus platform. Four flaps were added on two sides of the skull for fixation screw installation.

**Fig. 3.**
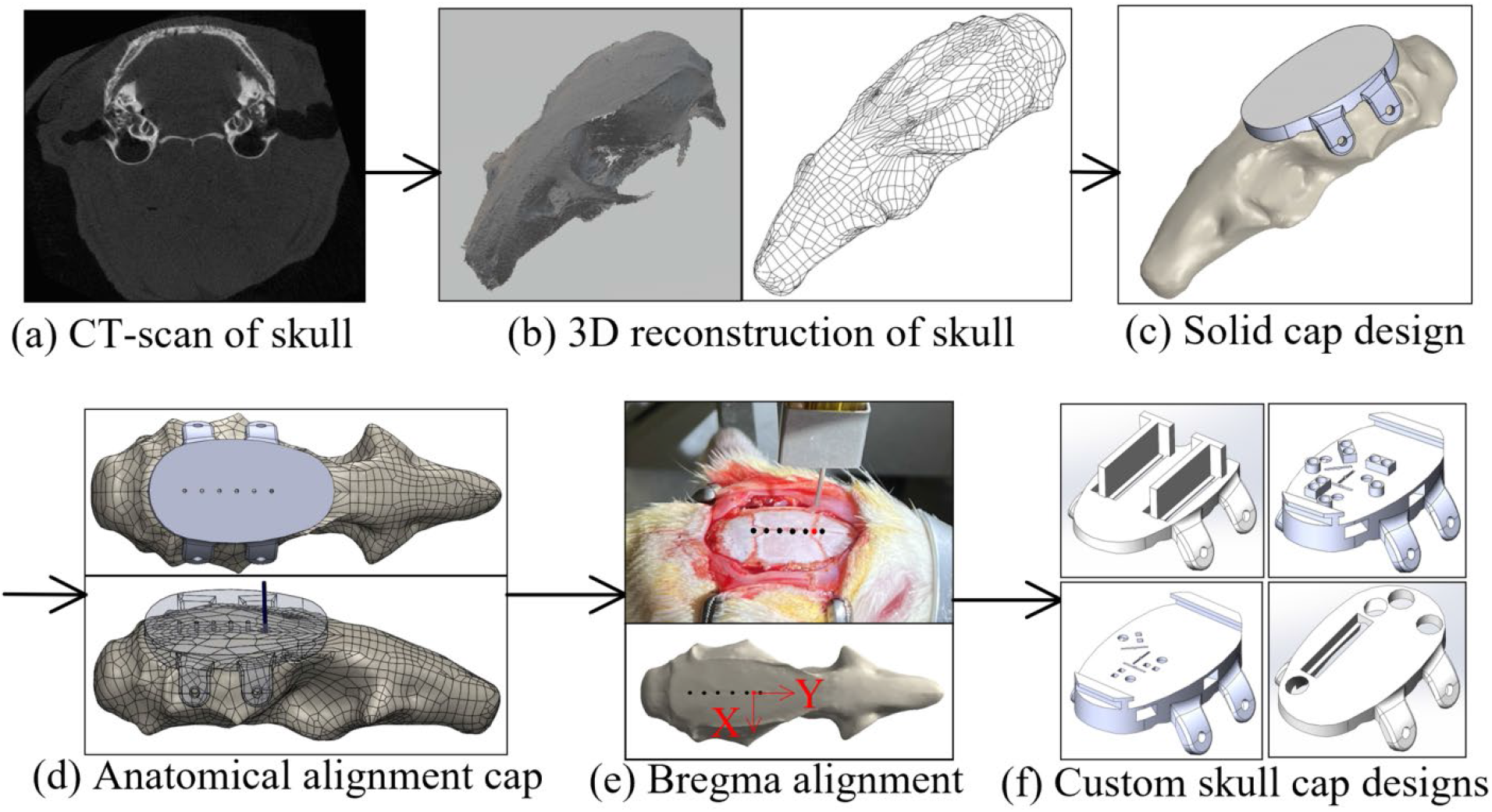
Design of the 3D printed skull cap and alignment with the anatomical feature.

To ensure a universal headcap design was capable of accurately representing any Sprague-Dawley’s true bregma location, we aimed to create a universal reference point on the headcap to serve as the bregma reference location. To achieve this aim, we used a stereolithography (SLA) 3D printer to create an alignment test headcap with six (6) 0.5 mm diameter through-holes separated by 2.2 mm (Fig. 3(d)). We attached this reference headcap with screws onto Sprague Dawley rats (n=6) varying in age, size, and body weight (Table 1) (surgical steps to be elaborated in Section 2.4). On each rat, a 0.5 mm male metal pin header (Model a18040700ux0248 by Uxcell, Hong Kong, China) was lowered into each reference hole on the cap, and stereotaxic coordinates were documented. Without moving the animal’s position on the stereotax, the pin header was lifted above the animal, and the headcap was removed. The pin header was then lowered back down to the true bregma position on the animal’s skull, and the coordinates were again documented. We used these coordinates to calculate the X/Y distance from each of the reference holes to the true bregma location on the skull for each animal, as in Fig. 3(e). The average X/Y distance from all animals was used to determine the location of a universal bregma point on the headcap. This universal headcap bregma point (X/Y coordinate 0,0) was then used as the reference for all implant target sites for all headcaps. Examples of custom recording location designs based on universal bregma are shown in Fig.3(f). Precision and accuracy of universal bregma compared to each animal’s true bregma are detailed in the results section.

**Table 1.** Age and weight of all rats used in bregma localization and error trials.

| Rat | Age | Weight |
| --- | --- | --- |
| 1 | 6 weeks | 360g |
| 2 | 6 weeks | 360g |
| 3 | 4 weeks | 266g |
| 4 | 4 weeks | 265g |
| 5 | 5 months | 605g |
| 6 | 6 weeks | 405g |

For the proof-of-concept animal surgery in this study, features to be supported and/or guided by the custom skull cap included four (4) microwire MEAs and two ground screws. For each MEA, the skull should provide accurate positions and orientations along the medial/lateral and anterior/posterior directions as well as guidance and stop features along the implantation direction. As shown in Fig. 4, based on the Bregma aligned skull cap, the custom skull cap design introduced two half-circle openings on the back for ground screw positioning (Fig. 4(b)). A rectangular-shaped opening was cut at the insertion location of each MEA. Two T-shaped structures were created next to the openings so that during insertion, the back and side edges of the MEA PCB could slide firmly against the corresponding support structure edges (highlighted in red in Fig. 4(b)) without the need for manual stereotactic alignment along each direction. Within the insertion opening of each MEA, a thin (0.25 mm thickness) step was created as the insertion depth indicator, as highlighted in red in Fig. 4(c). At the end of each MEA’s insertion, the PCB could sit on the 3D printed step free of extra support needed until the structure is fixed with dental cement. Note that the exact location of the step (insertion stopper) was determined by the targeted brain structure depth, microwire overhang length from the MEA PCB, and skull cap thickness. All caps in this study were printed using the same SLA 3D printer (Form 3B by Formlabs Inc., Somerville, MA) and material (Clear Resin V4 by Formlabs Inc., Somerville, MA) at a 25μm resolution setting. Post-processing was conducted for optimal part accuracy and performance (Form Wash and Form Cure by Formlabs Inc., Somerville, MA).

**Fig. 4.**
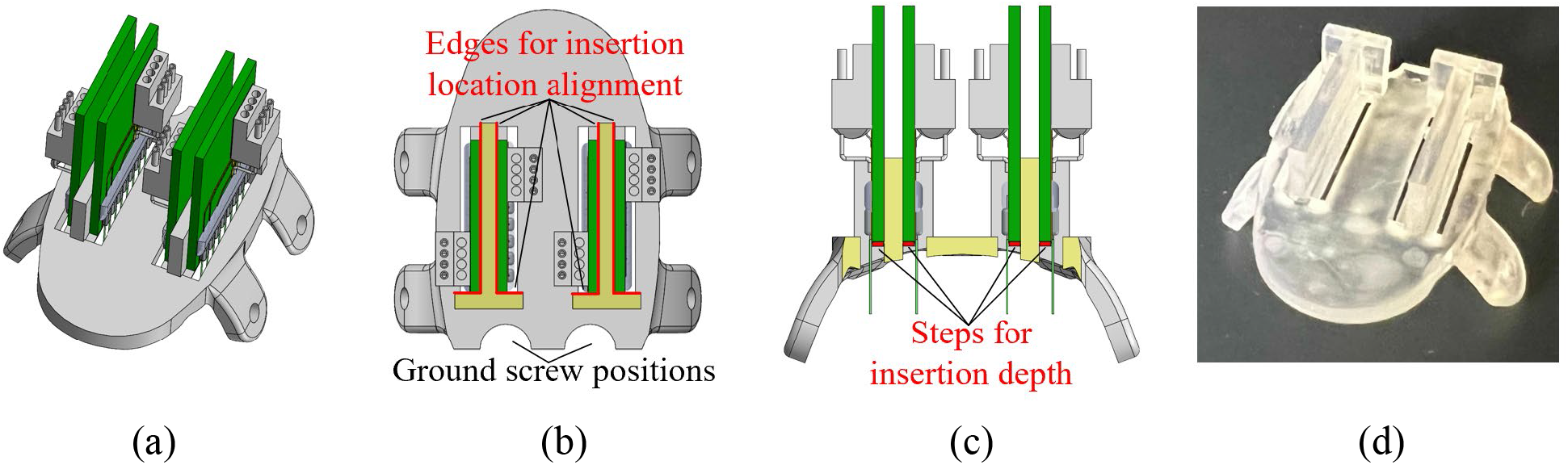
The custom skull cap used in this study: (a) design overview with four MEAs, (b) top and (c) front cross-sectional view of the cap showing position alignment feature designs, and (d) picture of the 3D printed skull cap.

As shown in Fig. 3(f), based on specific experimental needs, the thickness and overall size of the cap could be adjusted accordingly. If it is preferred to have the cap fulfill the craniotomy openings in the skull and rest against the brain membranes upon attachment, a 1.5 mm offset at the skull cap bottom was found in our preliminary study to fit most rats [26]. Support features and guidance openings could be designed on top of the cap, inside the cap, or offset beneath the cap bottom, based on the unique requirements of each experiment. This study presented an anatomical alignment with regard to rat Bregma, similar methodology could be applied to various species and anatomical structures.

### 2.3. Benchtop fabrication of microwire-based MEA

To fabricate the microwire-based MEAs used in this study, 50μm diameter tungsten microwires with polyimide insulation (CFW2029287 by California Fine Wires Co. Grover Beach, CA) were used as the raw material. The wires were hand-cut with fine surgical scissors (Artman Instruments, Kennesaw, GA) into 6 mm long segments and a 2.2 mm long insulation layer was stripped from each segment by a razor blade (Fig. 5(a)). To align multiple microwires with the gold pads on the MEA PCB, an alignment fixture was designed and 3D printed (Fig. 5(b)). The fixture had seven (7) evenly spaced ridges, forming eight (8) trenches whose centerlines coincident with the eight PCB gold pads (Fig. 5(c)). The width of each trench (0.6 mm in this study) was chosen as a compromise between wire placement difficulty and parallel misalignment tolerance between wires in an array. The MEA PCB was first placed in the designed opening to align the gold pads with trenches. Stripped tungsten microwire was then placed in the middle of each trench with the stripped section laid over the gold pad (1.6 by 0.8 mm in size). The stripped insulation edge was aligned with the PCB edge so that exactly 3.8 mm overhang of each wire could be achieved. This overhang length, combined with the rest step in the skull cap design for the PCB and the removed skull cap thickness, would yield a desired 1.6 mm insertion depth into the rat brain as the targeted recording location (Fig. 5(c)). These wire handling and skull cap design parameters can be adjusted to change insertion depth. Once all the wires are aligned in the trenches, about 0.5 mm of the stripped tungsten wires are extruded beyond the top of the gold pad (Fig. 5(c)). A small portion of quick setting epoxy (ClearWeld by J-B Weld Company, Sulphur Springs, TX) was applied on this stripped wire section, as in Fig. 5(d). This epoxy strip along the back side of the gold pads not only held all microwires firmly in place on the PCB but also created insulation for any exposed tungsten tips.

**Fig. 5.**
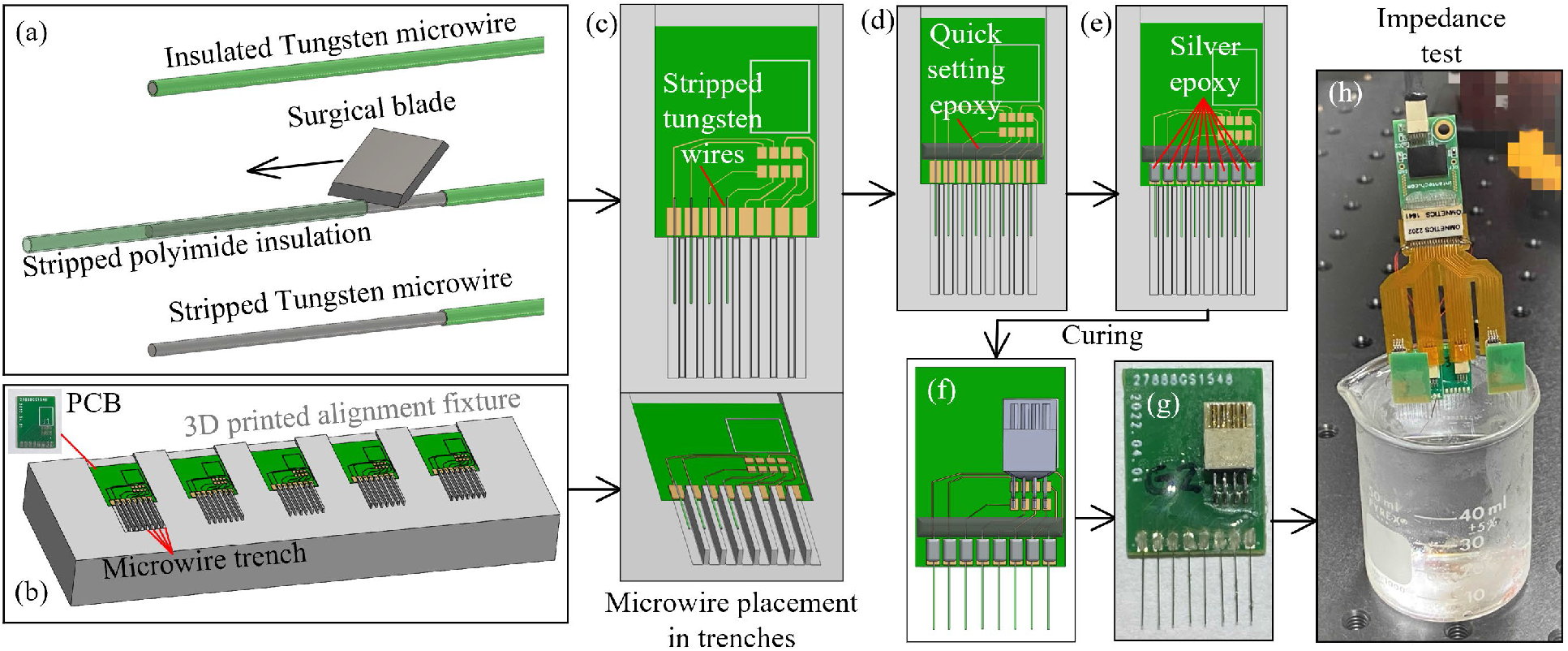
Fabrication process of the microwire-based MEA: (a) microwire insulation layer stripping with a surgical blade, (b) 3D printed fixture for microwire alignment, (c) microwire placement along the 3D printed trenches, (d) epoxy fixation microwires, (e) microwire electrical connection with silver epoxy, (f) and (g) connector soldering, and (h) impedance test.

Besides handling and placement, the soldering of the microwires was another labor-intensive and time-consuming step during MEA fabrication. Worse yet, some microwire materials like stainless steel and tungsten had nonoptimal solderability and would usually require extra steps of surface cleaning, pre-plating, pre-tinning, or specialized solder [18, 27, 28]. To overcome these challenges, conductive silver epoxy (EPO-TEK H20E by Epoxy Technology Inc., Billerica, MA) was applied using a sewing needle to create electrical connections between microwires and the PCB gold pads (Fig. 5(e)). The PCB with wires attached was heated in an oven at 150°C for one hour for the silver epoxy curing. Standard (non-silver) epoxy is then used to coat and insulate the proximal ends of the wires and pads to prevent any crosstalk. The 8-pin connector (A79613-001 by Omnetics Connector Corporation, Minneapolis, MN) was then soldered onto the rigid PCB board to finish the MEA fabrication process (Figs. 5(f) and (g)). Before implantation, each MEA went through an impedance test with its tips submerged into 1× phosphate-buffered saline (PBS), as shown in Fig. 5(h). Wire impedances were tested using a 32-channel headstage (RHD2132 by Intan Technologies, Los Angeles, CA) connected to a USB interface board (RHD2000 by Intan Technologies, Los Angeles, CA). All implanted MEAs had wire impedances of 20-200 kOhm measured at 1 kHz.

This modular assembly approach, 3D printed alignment fixture, and usage of silver epoxy could significantly lower the time and effort for MEA fabrication. For an inexperienced operator to build four 8-channel MEAs, it took about 3-4 hours including about 1.5 hours of wire trimming, 1.5 hours of wire alignment/fixation and connector soldering, and 1-hour idle time for silver epoxy curing. With an experienced operator, the entire process could be completed within 2-2.5 hours including the 1-hour oven curing time. This stands in significant contrast to often-expensive commercial solutions.

To generalize this benchtop MEA assembly approach, depending on fabrication resources available, the 3D printed alignment fixture with ridges could also be made by precision milling of metal blocks or chemical etching on silicon, to fulfill different MEA pitch size needs. If the raw microwire material was un-insulated when initially obtained, an extra insulation step (e.g., conformal coating) would need to be added at the beginning or after assembly. If the impedance test yielded a higher value than preferred, additional conductive coating (e.g., PEDOT:pTS, platinum-iridium) could be applied through electroplating [29]. MEAs with microwires in equal length were demonstrated in this study. The alignment fixture method could be easily applied to more complicated linear MEA configurations with various electrode lengths in an array.

### 2.4. Skull cap based MEA implantation and electrophysiological recording

Figure 6 presents the surgical procedures for implanting the fabricated MEAs at pre-printed locations through the custom skull cap into a certain depth in the rat brain. All animal experiments in this study were carried out in accordance with the University of Michigan Institutional Animal Care and Use Committee (protocol number PRO00009818 approved for 7/13/2020-7/13/2023). Electrode insertions were performed on Sprague-Dawley rats (Charles River) 3-6 months old. All animals were housed on a normal light dark cycle (lights on 7am-7pm) in a standard enriched environment and fed ad libitum.

**Fig. 6.**
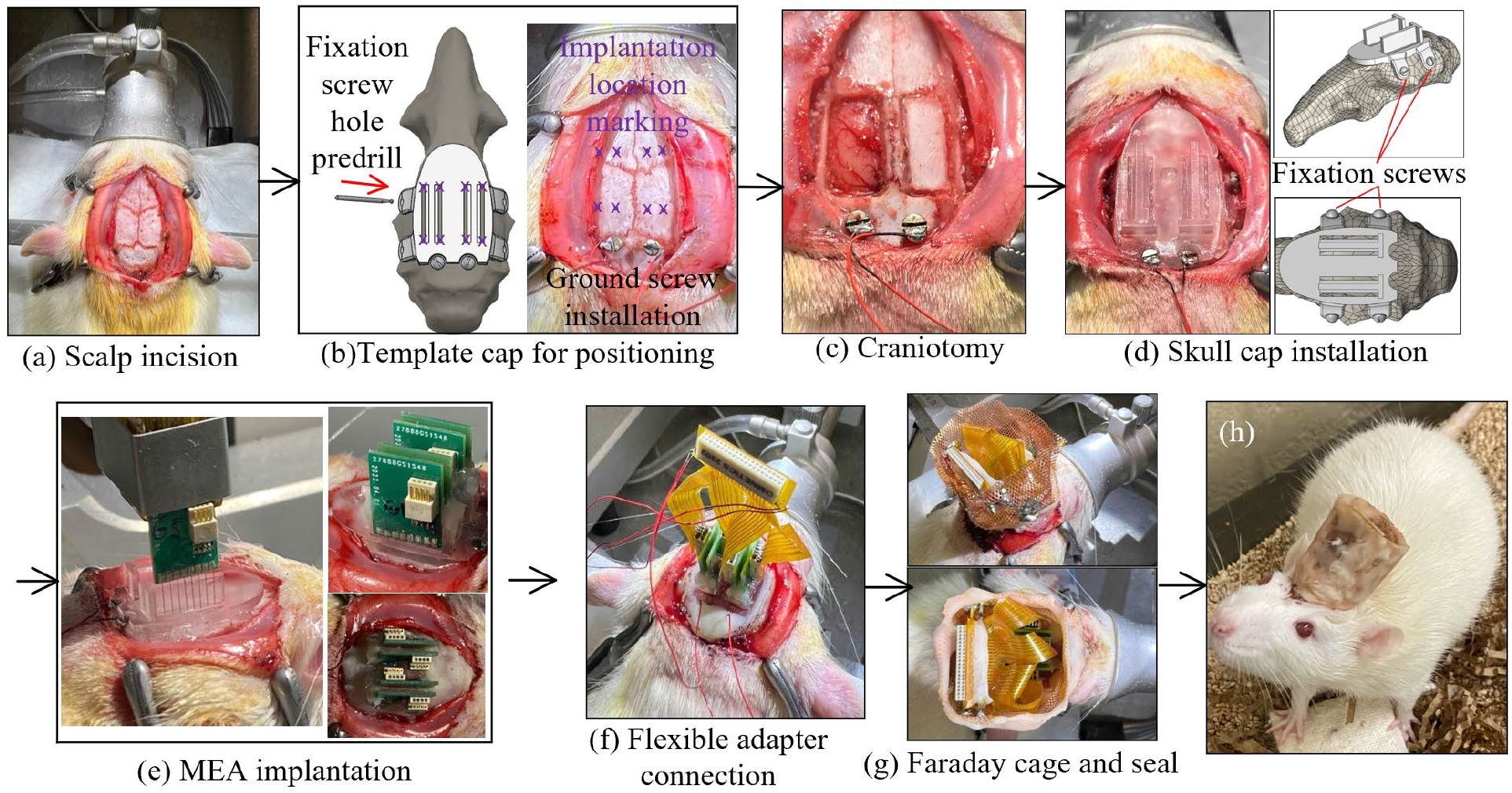
Surgical procedure for skull cap installation and cap-based MEA implantation.

Rats were anesthetized with isoflurane, placed in a stereotaxic apparatus, and given a subcutaneous injection of Carprofen 5mg/kg for analgesia as well as Methylprednisolone 30mg/kg intraperitoneally to limit brain swelling. A local anesthetic, Bupivacaine 1mg/kg, was administered subcutaneously to the scalp before surgical incision. An incision was made on the scalp along the midline of the skull with a sterile scalpel blade to expose the skull (Fig. 6(a)). Before surgery, a template skull cap was designed and printed (Fig. 6(b)). It had the same geometry as the final skull cap as in Fig. 4(d), except that it did not have the T-shaped support structure for easier reach of the insertion opening bottoms. This template cap was placed onto the skull surface and adjusted around for the best fit. It served as the surgical coordinate template and eliminated the needs for stereotactic measurements of all skull operation locations. A dental drill (MH-170 by Foredom, Bethel, CT) with the burr bit size of 0.9 mm diameter was used to pre-drill four fixation screw holes on the side of the skull for later anchoring the headcap to the skull. Two stainless steel screws (00-90, 3/32”) were driven through the skull over the cerebellum to serve as ground and reference. The burr was also inserted through the four MEA implantation openings and made marks on the skull as craniotomy reference locations (Fig. 6(b)). After template cap removal, two craniotomy windows were cut on the skull followed by durotomy of the opened regions (Fig. 6(c)). The craniotomy windows were filled with petroleum jelly, and the custom 3D printed skull cap as shown in Fig. 4(d) was then attached to the skull by tapping four fixation screws (00-90, 1/8”) through the pre-drilled holes (Fig. 6(d)). During attachment, both fixation screw holes and two installed ground screws served as the reference points to ensure proper alignment of the cap. Dental acrylic (UNIFAST Trad by GC America Inc., Alsip, IL) was used to seal the headcap-skull junction along the cap perimeter.

The MEA assembled as elaborated in Section 2.3 was gripped by a stereotactic arm (Model 1771 by David Kopf Instruments, Tujunga, CA) and aligned against the T-shaped structure on the custom skull cap as shown in Fig. 6(e). The MEA was then inserted until the PCB lowered into contact with the insertion depth indicator step as in Fig. 4(c). During the implantation, the stereotactic arm was used only as a manipulator and no manual quantitative distance measurement was needed. After implantation, as in Fig. 6(e), each MEA was anchored onto the skull cap with dental acrylic (UNIFAST Trad by GC America Inc., Alsip, IL). When all four MEAs were secured, the flexible adapter PCB was used to connect four 8-pin Omnetics connectors on MEAs to a 36-position connector for the headstage (Fig. 6(f)). To protect the recording apparatus from external electromagnetic waves, a Faraday cage was built around the PCBs with copper mesh and the ground screw wires were connected to the cage. The cage base was cemented to the perimeter of the headcap, and the cage body was covered with a layer of dental acrylic as shown in Figs. 6(g) and (h). The rats recovered for seven days, then electrophysiological recordings were performed in the rats’ home-cage using a 32-channel headstage (Intan Technologies RHD2132) connected to a USB interface board (Intan Technologies RHD2000) and sampled at 20kHz.

## 3. Results

### 3.1. Bregma Localization

To determine the precision and accuracy of universal headcap bregma and ensure that prefabricated headcaps have accurate insertion targets for any Sprague-Dawley rat, we measured the error in X/Y distance between universal headcap bregma and each animal’s true bregma location. The full range of error between true bregma and universal headcap bregma was within +/−1.0 mm along the A/P axis and +/−0.3 mm along the M/L axis (average A/P error distance +/−s.e.m. = 0.0 +/−0.25mm, average M/L error distance +/−s.e.m. = 0.0 +/−0.07mm, n=6; as in Fig.7). The precision of headcap positioning (using bregma as the reference) across six animals of varying age, size, and weight (Table 1) demonstrates the universality of the headcap design. The accuracy of this universal bregma point allows us to fabricate custom target openings on the cap prior to surgery and be able to reliably implant our arrays without stereotactic guidance.

**Fig. 7.**
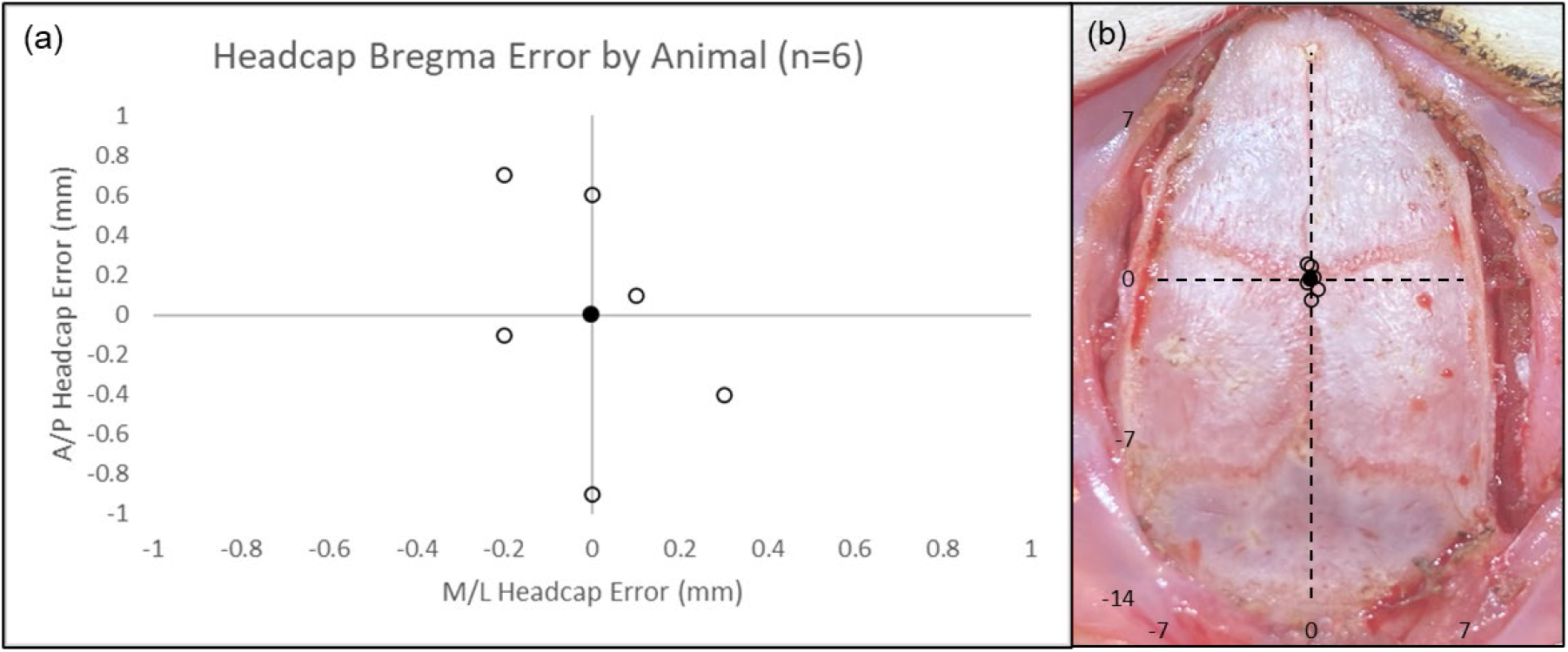
Headcap bregma location compared to true bregma of each animal: (a) Plot of difference between universal headcap bregma and each animal’s true bregma coordinates. Filled circle is average of 6 animals and represents the bregma coordinate (0,0) for the headcap. Open circles show distance (mm) from headcap bregma to each individual animal’s true bregma location. (b) Plot from (a) overlayed onto picture of typical dorsal skull (centered at bregma) to emphasize precision and accuracy of universal headcap bregma. Skull surface measures approximately 14×24 mm.

### 3.2. Brain recording results

Four MEAs containing 8 wires each were inserted through the headcap as detailed above. The 32 wires formed a rectangular grid spanning 7mm along the anterior/posterior axis and 10mm along the medial/lateral axis to allow for broad sampling of cortical activity. All wires were inserted between the coronal and lambdoid skull sutures to a depth of 1.6 mm. High-resolution recordings were collected at 20 kHz to obtain local field potential and single unit signals.

LFP from 32 channels was recorded over many weeks during home-cage sessions, each lasting 2-6 hours (Fig. 8(a)). Presence of delta oscillations (0.5-4 Hz) during non-rapid-eye-movement (NREM) sleep indicate normal physiology and successful cortical LFP recordings (Fig. 8(b)). Single units were also observed in multiple channels for more than 5 months (Fig. 8(c)), indicating stable high-resolution recording capabilities. Despite removing a large portion of the dorsal calvarium, none of the animals showed behavioral changes or deficits, and there were no signs of infection or other complications. Signals could be maintained for multiple months, in a manner comparable with more traditional surgery methods.

**Fig. 8.**
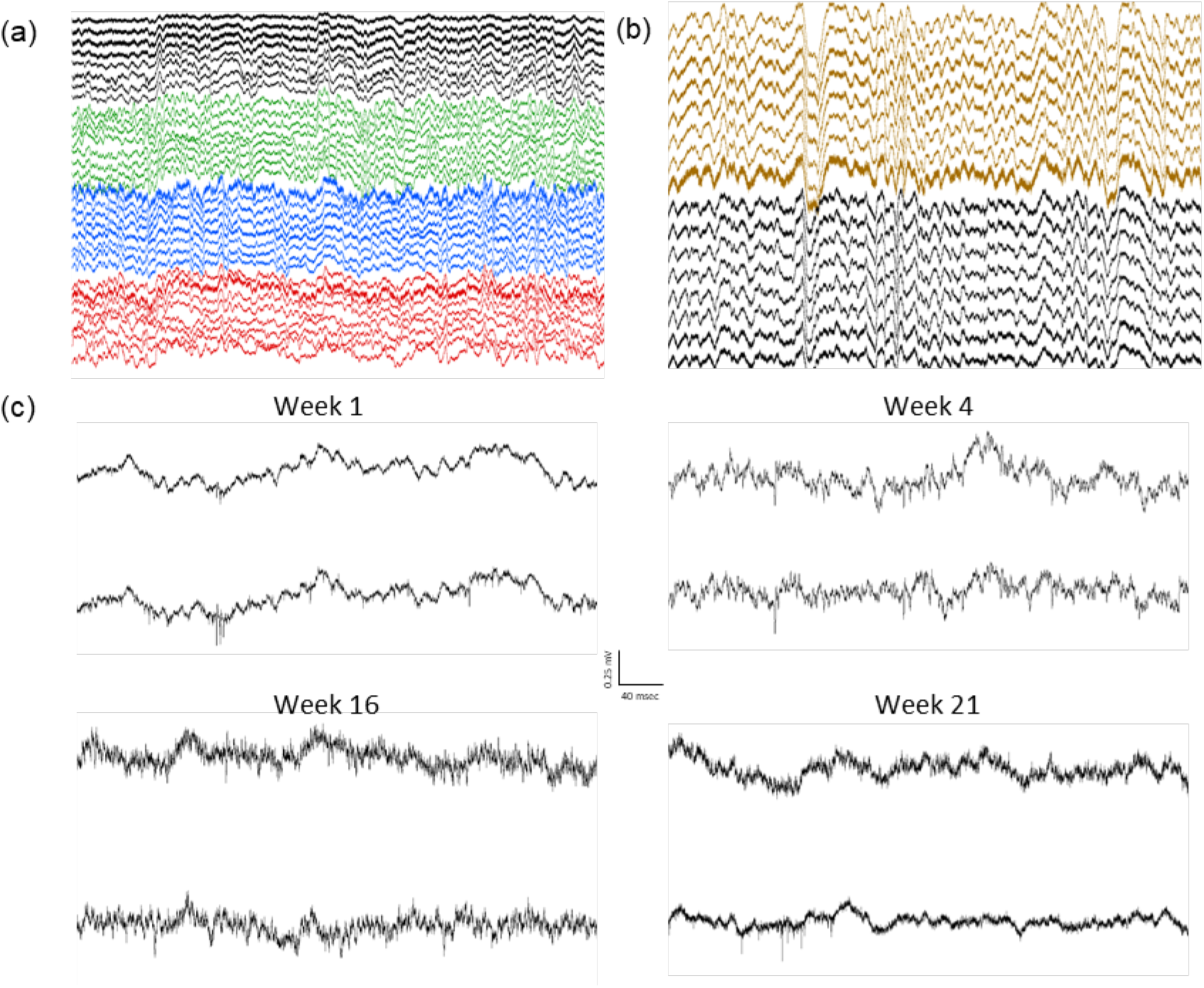
Example recordings from an animal with four 8-wire array implants: (a) Example two seconds of 32-channel recording. Individual arrays are highlighted in different colors. (b) Example four seconds showing delta oscillations (0.5-4 Hz) characteristic of normal physiology during NREM sleep. (c) Spiking activity in two example channels from the same animal over a 5-month period.

## 4. Discussion

Here we have shown the development of a new methodology based on a computer-aided designed and 3D printed skull cap system for both MEA fabrication and rat brain surgery. This work reduces labor and time for creation and surgical implantation of MEAs and allows laboratories to undertake fabrication on their own at low cost. The printed skull replacement with ability to pre-shape and pre-form craniotomies and mounts for surgeries provides a system that is convenient and effective now, but also is an important first step for future developments.

We have shown that the top portions of the skull can be replaced by printed parts without compromising animal health, behavior, or recording quality. We’ve also shown that our cap can be used to insert many wires that yield stable and high-resolution recordings over multiple months. Our CT-aided design conforms well to the skulls of Sprague-Dawley rats of varying size, age, and weight, and creates minimal bregma alignment error. For higher resolution alignment needs, CT head scan (bone) and T1 magnetic resonance imaging (MRI) scan (brain structure) could be co-registered to directly correlate custom skull cap design with targeted brain structures [26].

Most of the design, fabrication, and implantation methodologies [26] showcased in this proof-of-concept study could be generalized to electrophysiological recording and stimulation on different species with various devices and configurations, thanks to the capability provided by CAD and 3D printing. Because our caps are 3D-printable, they are affordable, widely accessible, and could also be tailored for custom recording locations for specific regions or more broadly for large-scale cortical recordings. This new implantation method could also accommodate other recording designs including micro-drive manipulators, tetrodes, silicon probes, optogenetic fibers, and imaging windows. Each of these recording options can be planned prior to surgery to reduce manual manipulation and implantation difficulty, resulting in surgeries that are more efficient and repeatable, offer high-resolution and high-density and/or broad-scale sampling, and provide stable chronic recordings.

Given that brain dynamics involve multi-region coordination, the ability to better study those dynamics will depend on the development of systems to implant complex arrays of probes that are not practically feasible with manual methods. Computer-based design and printing of surgery-enabling platforms is very likely to become increasingly commonplace as rodent brain surgeries become more complex. Our system represents an important step towards computer-aided rodent brain implantation to better study the brain and disease models from the perspective of broad-scale electrophysiologic dynamics.

## 5. Conclusions

A new methodology for laboratories to fabricate low-cost custom microwire-based MEAs and implant them at custom multiple brain regions based a computer-aided designed and 3D printed skull cap platform has been developed. The skull cap design based on CT-scan of a single Sprague-Dawley rat conforms well to the skulls of Sprague-Dawley rats of varying size, age, and weight, and creates minimal bregma alignment error. Based on the proof-of-concept design, prototype, and animal recording study, the easily duplicable recording system was shown capable of recording spiking activities over multiple months. The design, fabrication, and implantation methodologies could be easily reconfigured and generalized to various devices and configurations on different species, to enable low-cost highly custom multi-region recording/stimulation studies.

## CRediT authorship contribution statement

**Dongyang Yi**: Conceptualization, Methodology, Validation, Investigation, Writing - Original Draft, Visualization. **Jeremiah P. Hartner**: Conceptualization, Methodology, Validation, Investigation, Data Curation, Writing - Review & Editing, Visualization. **Brian S. Ung**: Investigation, Writing - Review & Editing. **Harrison L. Zhu**: Investigation, Writing - Review & Editing. **Brendon O. Watson**: Conceptualization, Methodology, Writing - Review & Editing, Supervision, Project administration, Funding acquisition. **Lei Chen**: Conceptualization, Methodology, Writing - Review & Editing, Visualization, Supervision, Project administration, Funding acquisition.

## Acknowledgements

This work was supported by the National Institutes of Health (R21 MH120465).

